# Rhes Deletion Prevents Age-Dependent Selective Motor Deficits and Reduces Phosphorylation of S6K in Huntington Disease Hdh150Q(CAG) Knock-In Mice

**DOI:** 10.1101/2021.06.16.448681

**Authors:** Jennifer Hernandez, Neelam Shahani, Supriya Swarnkar, Srinivasa Subramaniam

## Abstract

Huntington disease (HD) is a neurodegenerative disease caused by a CAG trinucleotide repeat expansion in the huntingtin (mHTT) protein. This expansion is thought to promote striatal atrophy by a combination of cell- and non-cell-autonomous processes, but the mechanisms are unclear. Previous evidence suggests that the striatal-enriched SUMO E3-like protein Rhes could play a pathological role in HD. Rhes interacts with, and SUMOylates, mHTT and promotes toxicity and Rhes deletion ameliorates the HD phenotype in cell and severe mouse models of HD. However, the effect of Rhes on less severe knock-in models of HD remains obscure. Here, we report that a Hdh(CAG)150 knock-in murine model of HD showed diminished body weight but no changes in locomotor coordination or activity at 80 and 100 weeks of age. Conversely, Rhes deletion did not impact the body weight or behaviors but caused a significant reduction of gait, clasping, and tremor behaviors in Hdh^150Q/150Q^ mice. Rhes deletion did not affect the loss of striatal DARPP-32 protein levels but abrogated the hyper ribosomal protein S6 kinase beta-1 (S6K) phosphorylation, which is a substrate for a mechanistic target of rapamycin complex 1 (mTORC1) signaling, in Hdh(CAG)150 mice. Interestingly, striatal Rhes protein levels were downregulated in the striatum of Hdh(CAG)150 mice, indicating a potential compensatory mechanisms at work. Thus, Rhes deletion prevents age-dependent behavioral deficits and diminishes hyperactive mTORC1-S6K signaling in Hdh(CAG)150 knock-in mice HD striatum.

## Introduction

Wild-type huntingtin (wtHTT) is a ubiquitously expressed protein with a polyglutamine tract encoded by the *HTT* CAG repeat expansion (mHTT) that causes Huntington Disease (HD). HD is characterized by the early loss of medium spiny neurons (MSNs) in the striatum, and this loss subsequently affects motor and cognitive functions^[1, 2]^. As HD progresses, it affects other parts of the brain, as well as the peripheral tissues^[3, 4]^. The mHTT protein and its proteolytically cleaved fragments affect several cell functions, such as vesicle- and microtubule-associated protein/organelle transport, calcium dysregulation, transcription, autophagy, and sphingosine and cysteine metabolism, as well as having effects on tissue maintenance, secretory pathways, and cell division^[5-22]^. However, the mechanisms underlying HD pathogenesis remain unclear, and the currently available drugs, although providing symptomatic relief, neither prevent nor slow HD progression. Elucidating the mechanism governing striatal abnormalities should therefore provide opportunities for the development of novel and effective therapeutic interventions.

One possible target for these interventions is Rhes, which is highly expressed in the dopamine 1 receptor (D1R), D2R MSNs, and cholinergic interneurons in the striatum^[23, 24]^. It is also expressed to some extent in other brain regions, such as the cortex and hippocampus^[23, 25, 26]^. Rhes expression is induced by thyroid hormones, and Rhes can inhibit the cAMP/PKA pathway and N-type Ca^2+^ channels (Cav 2.2)^[24, 27-29]^. We have found several new roles for Rhes in the striatum; for example, Rhes can bind directly to and activate mTOR in a GTP-dependent manner, thereby promoting L-DOPA-induced dyskinesia (LID) in a pre-clinical model of Parkinson’s disease,^[30]^ in agreement with a recent report^[31]^. We also found that Rhes harbors a C-terminal SUMO E3-like domain and physiologically regulates SUMO modification via the cross-SUMOylation of the E1 (Aos1) and E2 (Ubc9) enzymes^[32]^. We recently identified several novel nuclear SUMO substrates of Rhes and a role for Rhes in gene regulation^[33]^.

We have demonstrated that Rhes-SUMO signaling plays major roles in HD. For example, we found that Rhes interacts with mHTT and promotes cellular toxicity by increasing the soluble forms of mHTT via SUMO1 modification^[34]^, in agreement with an independent study^[35]^. The toxicity of Rhes expression in HD has also been independently demonstrated in various cell and hESC-derived models of HD^[36-40]^, but the underlying molecular mechanisms of Rhes-mediated toxicity remain unclear. Interestingly, Rhes promotes tau toxicity by regulating lysosomal activity and SUMOylation in Tg4510 tauopathy mouse^[41]^, and altered localization and clearance of neuronal Rhes is considered as a novel hallmark of tauopathies^[42]^. Serendipitously, we found that Rhes can be robustly transported between cells and that it functions as a critical determinant of mHTT transport between neuronal cells via the tunneling nanotube-like (TNT-like) “Rhes tunnel” protrusion^[43, 44]^. The mHTT cargoes travel through this protrusion and along the membrane of acceptor cells before being internalized^[43]^. By contrast, SUMOylation-defective mHTT is transported much less efficiently, indicating the importance of SUMO in Rhes-mediated mHTT transport^[43]^. These results indicate that Rhes–SUMO signaling participates in HD pathogenesis.

Consistent with the above studies, we found that Rhes deletion ameliorated HD pathogenesis in a number of different transgenic (Tg) HD models. Rhes deletion had a marked effect on preventing motor phenotype in male, but not in female, N171-82Q HD Tg model mice^[45]^. *Rhes*^*–/–*^ also exhibits a gender effect on dopaminergic-drug induced motor behaviors^[24]^. A previous independent study showed that Rhes deletion in R6/2, a more severe HD model, also prevented HD-associated motor deficits^[46]^. However, we found that Rhes overexpression worsened the HD-related motor deficits accompanied by enhanced soluble forms of mHTT in the striatum of 6-month-old *Hdh*^*150Q/150Q*^ knock-in (KI) mice, which express full-length mutant huntingtin with slowly developing and less severe behavioral phenotype mouse^[47] [48]^, and in the cerebellum of N171-82Q mice^[45, 48]^. A recent study showed that HAP1 deficiency increases the binding of Rhes to soluble N-terminal mutant HTT and to SUMOylated N-terminal HTT, while promoting selective neuronal loss in the striatum of full-length mutant HTT KI (140CAG) mice^[49]^. These multiple independent results^[34-40, 45, 46, 49, 50]^ indicate that Rhes plays a major role in striatal neuronal loss by interacting with and modifying mHTT and that its role is modulated by additional proteins in the striatum.

By contrast, microRNA-based depletion of *Rhes* mRNA in the striatum of N171-82Q showed no protective effect. Instead, its depletion in 12-month-old BACHD mice (similar to N171-82Q;*Rhes*^*–/–*^)^[24]^ resulted in displays of mild hypoactivity, as well as a ∼4% decrease in striatal volume as measured by MRI^[51]^. Overexpression of AAV-Rhes in the striatum of N171-82Q mice improved their rotarod performance^[52]^; however, in that study^[52]^, Rhes was flag-tagged at the C-terminal end and this could have affected farnesylation and mislocalized Rhes to the nucleus^[34]^. Nevertheless, these results indicated that Rhes-mediated HD pathogenesis in mice in vivo depends on the choice of HD mouse model, as each may invoke as yet unknown compensatory mechanisms to modify the Rhes-mediated mHTT toxicity. However, the role of Rhes deletion in modulating HD pathogenesis in less severe and late-onset knock-in HD mouse models remains obscure.

Hyperactive mTORC1 is observed in HD by independent studies^[53-55]^, but its role in influencing HD remains controversial. For example, while rapamycin prevented HD phenotype^[55]^, the overexpression of Rheb, an activator of mTORC1, improved HD-like symptoms in mice^[52]^. In contrast, Rheb overexpression in Drosophila worsened HD-like phenotype^[56]^. We found depleting TSC1 exacerbated mTORC1 in the striatum and worsened HD-like phenotype in mice^[55]^. Thus, mTORC1 differentially affects HD, depending upon HD models and assays used to interpret the phenotype. In addition, mTORC1 activation, a key metabolic hub that is controlled by various upstream stimuli may be regulated by mHTT in cell/tissue-dependent manner.

In the present study, we further examined the in vivo role of Rhes on late-onset HD using a Hdh150Q(CAG) KI mouse model^[47, 48]^. We crossed Rhes-KO with Hdh150Q(CAG) KI mice and carried out the motor deficit and biochemical parameter studies in both male and female mice.

## Results

### Behavioral assessment of Rhes deletion in Hdh^150Q^ knock-in mice at 80 weeks

Constitutive depletion of Rhes (Rhes-KO) in the Tg HD R6/2^[46]^ and N171-82Q ^[45]^ mice prevented HD-related behavioral and pathological changes. However, the effect of a Rhes-KO in knock-in HD mouse models remains unknown. To test this question, we crossed Rhes-KO mice with Hdh150Q(CAG) KI and generated Rhes deleted *Hdh*^*150Q/150Q*^ homozygous and *Hdh*^*150Q/+*^ heterozygous mice, which developed a late-onset age-dependent HD, with subtle but significant motor abnormalities appearing between 70 and 100 weeks of age^[47, 48]^. The number of animals (mixed sex ratio) used is indicated in Table 1.

**Table 1.** Number of animals used

| **Table 1. Number of animals used** |  |  |
| --- | --- | --- |
| **Genotype** | **80 weeks** | **100 weeks** |
| Rhes WT/HD-het | 19 | 12 |
|  | (11 male) | (7 male) |
| Rhes KO/HD-het | 14 | 12 |
|  | (7 male) | (7 male) |
| Rhes KO/HD-homo | 10 | 7 |
|  | (5 male) | (2 male) |
| Rhes KO/WT | 12 | 9 |
|  | (5 male) | (3 male) |
| Rhes WT/HD-homo | 10 | 6 |
|  | (3 male) | (1 male) |
| WT | 9 | 5 |
|  | (4 male) | (2 male) |

At 80 weeks, no significant effect was observed on body weight in the *Hdh*^*150Q/+*^ mouse groups compared to the control groups (Fig. 1A), consistent with previous reports^[47, 57, 58]^. The *Hdh*^*150Q/150Q*^ mice (n = 10; 3 male) showed a significant loss of body weight, consistent with a previous report^[47]^. This weight loss was similar to that seen in *Rhes*^*–/–*^;*Hdh*^*150Q/150Q*^ (n = 10; 5 male) mice compared to WT (n = 9; 4 male) and *Rhes*^*–/–*^ mice (n = 12; 5 male, Fig. 1A).

**Figure 1.**
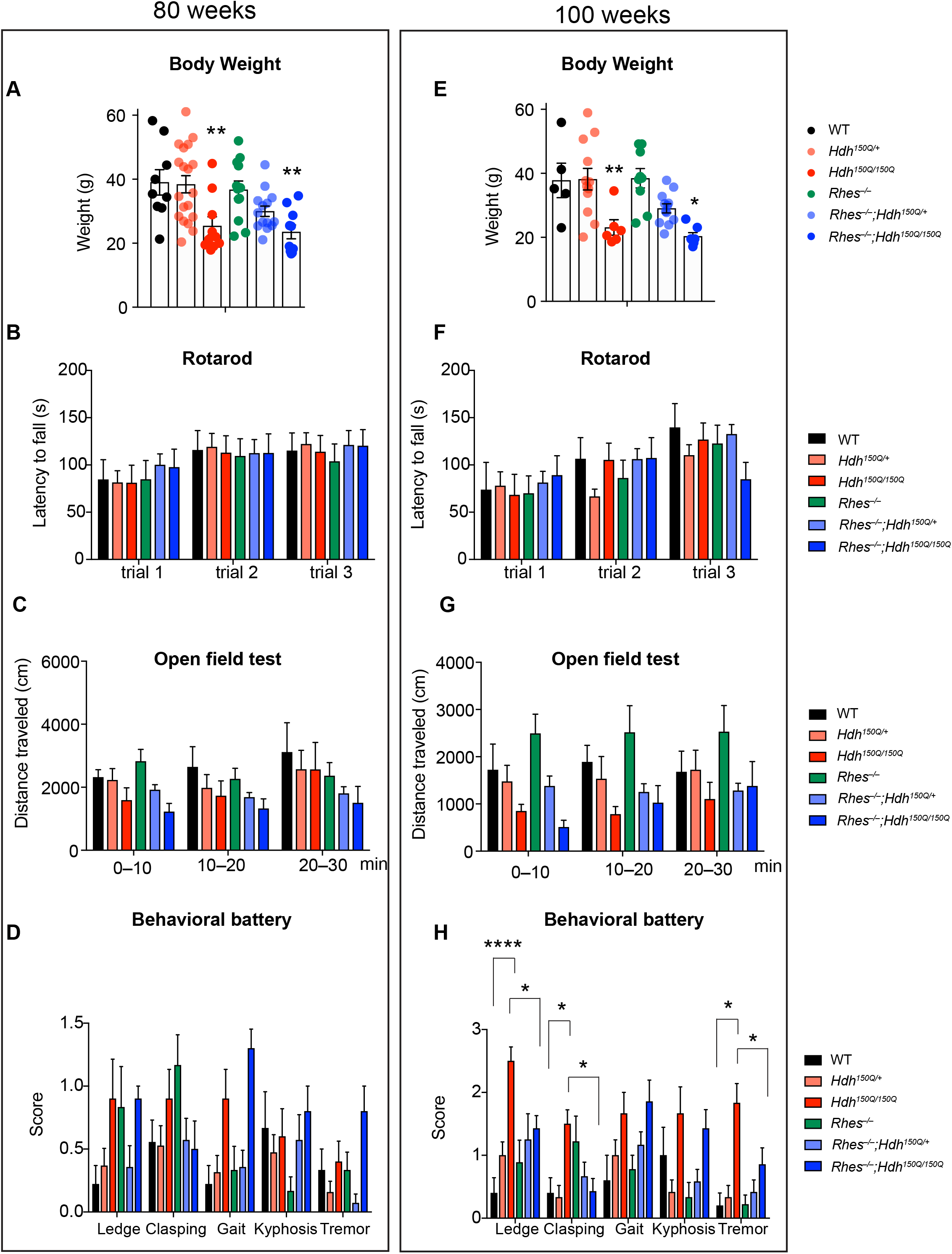
Effect of Rhes deletion on age-associated motor abnormalities in Hdh150Q KI HD mice. (A) Body weight, (B), rotarod, (C) open field (D) behavioral battery composite score included ledge-test, clasping, gait, kyphosis, and tremor evaluation. The frequency of each behavior according to the genotype is shown in F, a composite score for all the behavioral battery is shown in G. Data are mean ± SEM (n = 5-19/genotype). *P<0.05, **P<0.01, ****P< 0.0001, 2way ANOVA followed by Tukey’s multiple comparison test.

The rotarod behavior tests conducted at 80 weeks in *Hdh*^*150Q/+*^ or *Hdh*^*150Q/150Q*^ mice did not reveal any significant defect in the locomotor coordination, and the performance in three different trials was comparable to that of the WT, *Rhes*^*–/–*^;*Hdh*^*150Q/+*^ or *Rhes*^*–/–*^ ;*Hdh*^*150Q/150Q*^ groups (Fig. 1B), again consistent with previous studies^[47, 58]^. Two-way ANOVA of the three trials for each animal revealed no significant difference between the genotypes [F (2, 78) = 2.067, P=0.1334] or interactions [F (4, 78) = 0.06588. P = 0.9919]. Open field tests conducted at 80 weeks in the *Hdh*^*150Q/150Q*^ mice showed a decreasing trend in spontaneous locomotion in the *Hdh*^*150Q/150Q*^ mice, and a significant correlation was found between genotypes (F (2, 78) = 4.202, P = 0.0185), but not with the interactions (F (4, 78) = 0.1368, P = 0.9682) or time (F (2, 78) = 1.18, P = 0.3127).

Similarly, behavioral battery tests (ledge, clasping, gait, kyphosis, and tremor) conducted at 80 weeks revealed a significant genotype effect (F (2, 130) = 6.466, P=0.0021), but no significant interaction (F (8, 130) = 1.665, P=0.1128) or time (F (4, 130) = 0.7863). The *Hdh*^*150Q/150Q*^ mice and showed a trend toward significant defects in the ledge (P = 0.0595) and gait tests (P = 0.0595), and the *Rhes*^*–/–*^;*Hdh*^*150Q/150Q*^ showed significant defects in gait compared to WT (P = 0.0011).

### Behavioral assessment of Rhes deletion in Hdh^150Q^ knock-in mice at 100 weeks

By the age of 100 weeks, some mice (both male and female) died during the study in all groups: four out of nine (4/9) in the WT, 4/10 in the HD-homo, 3/12 in the Rhes-KO, 3/12 in the Rhes-KO;HD, 2/14 in the Rhes-KO;HD-het, and 7/19 in the HD-het groups.

Body weight remained lower for the *Hdh*^*150Q/150Q*^ mice but not in the *Hdh*^*150Q/+*^ groups, while Rhes deletion had no impact on body weight (Fig. 1E).

Rotarod tests showed no significant defects, even at 100 weeks between groups (Fig. 1F), even though previous studies had showed a slight deficiency in rotarod performance around this age in *Hdh*^*150Q/150Q*^ mice^[47, 58]^. The difference could reflect the different methodological parameters used. In the original study, no training was used, and the test was conducted at a fixed low-speed setting or with the accelerating rod method run for a maximum of 5 min. In the present case, at both 80 and 100 weeks, we performed the rotarod test using a linearly accelerating rotation paradigm in three trials separated by 20 min for four consecutive days, as described in our previous work^[55]^.

The open field test results showed a significant correlation between genotypes (F(2, 45) = 4.644, P = 0.0147) but no difference in the interaction (F (4, 45) = 0.511, P = 0.7279) or time (F (2, 45) = 0.7255, P = 0.4897) (Fig. 1G).

In the behavioral battery tests, the *Hdh*^*150Q/150Q*^ mice showed severe deficits in the ledge (P<0.0001), clasping (P<0.0388), gait (P < 0.0388), and tremor (P <0.0012) tests when compared to WT mice. Interestingly, the *Rhes*^*–/–*^;*Hdh*^*150Q/150Q*^ mice showed significantly decreased levels of ledge (P < 0.0266), clasping (P <0.0266), and tremor (P <0.0478) defects (Fig. 1H).

Altogether, the behavioral data indicated that *Hdh*^*150Q/150Q*^ mice exhibited late-onset but subtle motor deficits and that Rhes deletion had a significant rescue effect on selected neurological abnormalities at 100 weeks.

### Biochemical assessment of Rhes deletion in Hdh^150Q^ mice

After the behavioral tests, the heterozygous (*Hdh*^*150Q/+*^, Fig. 2) and homozygous mice (*Hdh*^*150Q/150Q*^, Fig. 3) were sacrificed, and their brains were removed for western blot analysis. We found decreased expression of DARPP-32, a striatal neuron marker, in *Hdh*^*150Q/+*^ and Rhes deletion did not impact DARPP-32 levels in *Rhes*^*–/–*^;*Hdh*^*150Q/+*^ (Fig. 2A, B).

**Figure 2:**
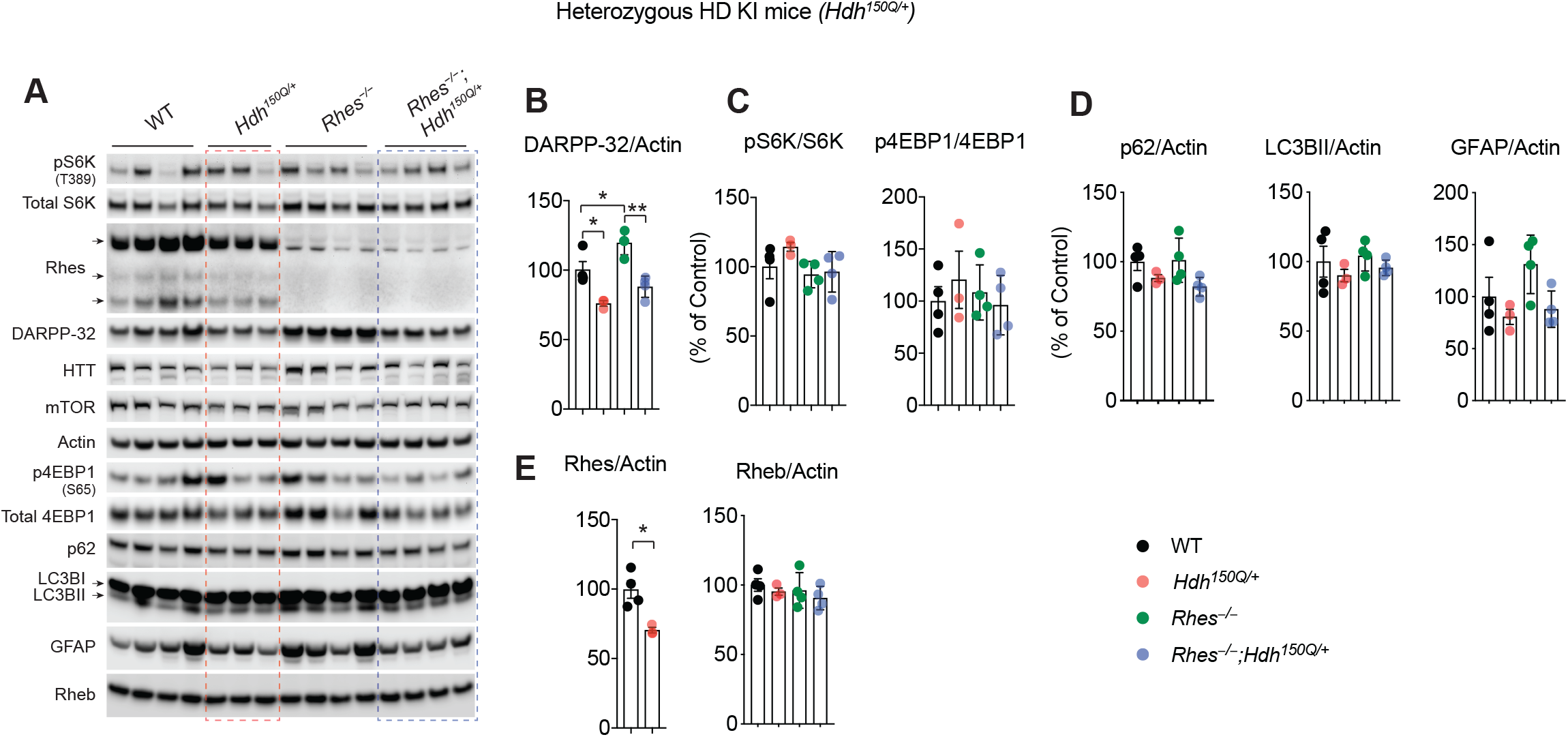
Biochemical alterations of Rhes deletion in 103 weeks old Hdh150Q KI heterozygous mice. (A) Western blot analysis of indicated proteins from striatum of indicated mice in Hdh^150Q/+^ heterozygous and other mice groups. (B-E) Bar graph shows quantification of the indicated proteins from E. Data are mean ± SEM, n = 3-4 per group, *P<0.05, **P<0.01, one-way ANOVA followed by Tukey’s multiple comparison test.

**Figure 3:**
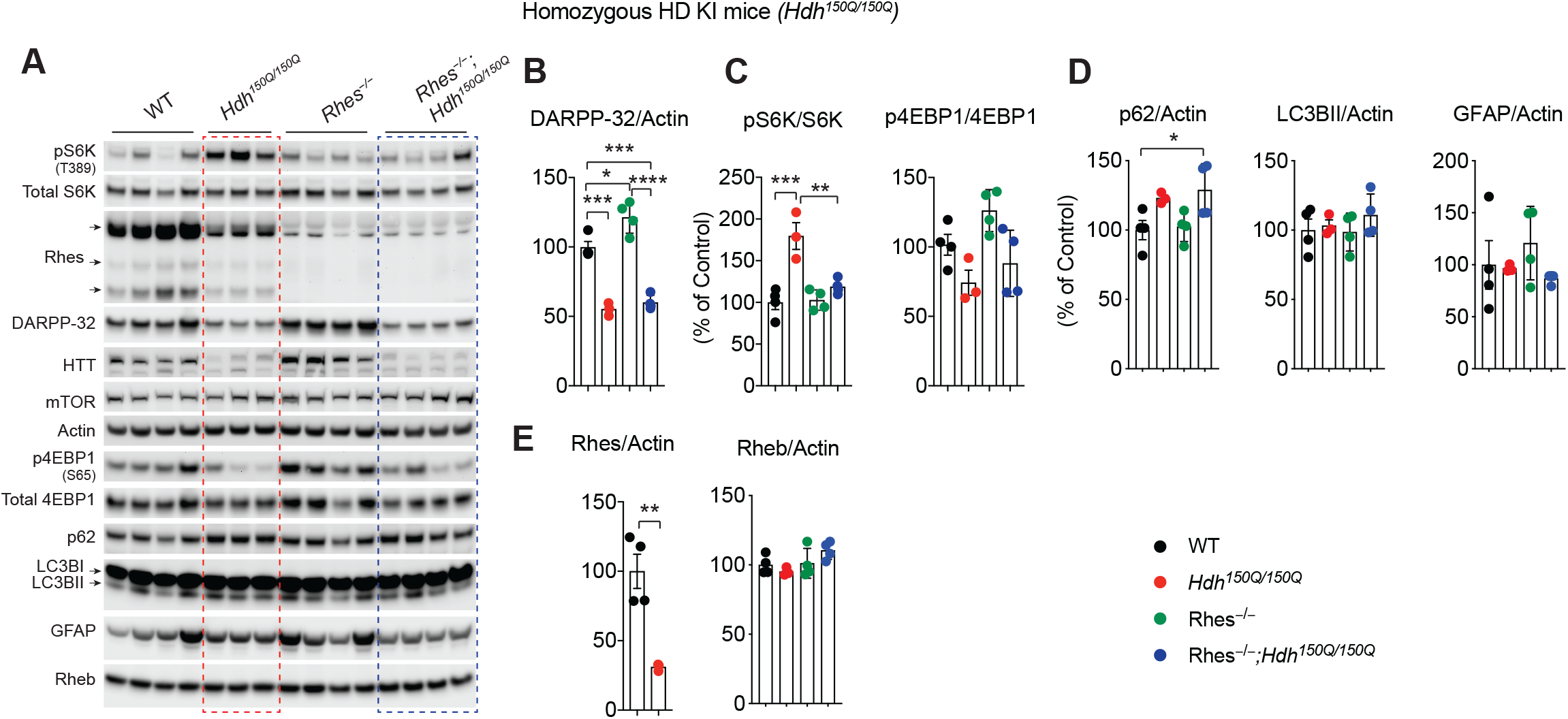
Biochemical alterations of Rhes deletion in 103 weeks old Hdh150Q KI homozygous mice. (A) Western blot analysis of indicated proteins from striatum of indicated mice in Hdh^150Q/150Q^ homozygous and other mice group. (B-E) Bar graph shows quantification of the indicated proteins from E. Data are mean ± SEM, n = 3-4 per group, *P<0.05, **P<0.01, ***P< 0.001, ****P< 0.0001, one-way ANOVA followed by Tukey’s multiple comparison test.

We previously showed that mHTT could elicit mTORC1 signaling in the HD cell model^[55]^, so we evaluated mTORC1 signaling in HD mice. We found that striatal mTORC1 signaling targets the phosphorylation of S6K (Thr389) or phosphorylated 4E-BP1 (Ser65) was not significantly upregulated in the striatum in *Hdh*^*150Q/+*^ mice (Fig. 2A, C).

Autophagy and inflammatory process are also altered in HD^[59, 60]^. So, we evaluated p62 and LC3BII (markers of autophagy) and GFAP (glial marker) and found no significant alteration of these markers in *Hdh*^*150Q/+*^ mice striatum by western blotting (Fig. 2A, D).

Interestingly, we found significant reductions in the Rhes levels, but not Rheb, another member of small GTPase^[61]^, in the *Hdh*^*150Q/+*^ mouse striatum (Fig. 2E).

In homozygous mice (*Hdh*^*150Q/150Q*^), the DARPP-32 levels were further strongly diminished in the striatum and Rhes deletion had no impact in *Rhes*^*–/–*^;*Hdh*^*150Q/150Q*^ mice (Fig. 3A, B).

The hyperphosphorylation of S6K (Thr389) was observed in the striatum of homozygous HD mice. Rhes deletion completely abolished this without affecting the basal pS6K, indicating that Rhes may promote mTORC1 signaling in the striatum under pathological conditions, consistent with previous work^[30, 31]^. Phosphorylated 4E-BP1 (Ser65), which in some instances can be mTORC1-independent^[62]^, however, is not significantly altered in the striatum of *Hdh*^*150Q/150Q*^ mice (Fig. 3A, B). Thus mTORC1–S6K pathway is particularly regulated by Rhes in *Hdh*^*150Q/150Q*^ mice.

Investigation into the autophagy regulators revealed while p62 levels show a trend of an enhancement, but the steady-state levels of LC3BII are unaltered in *Hdh*^*150Q/150Q*^ mice. Rhes deletion has no impact on p62 or LC3BII levels in *Rhes*^*–/–*^; *Hdh*^*150Q/150Q*^ mice (Fig. 3A, D).

Similar to heterozygous *Hdh*^*150Q/+*^ mouse striatum, a further dramatic reduction of Rhes in the *Hdh*^*150Q/150Q*^ homozygous HD KI striatum was observed, while the Rheb levels remain unaltered. (Figs. 3A, E).

Altogether, the biochemical results indicated that DARPP32 levels are diminished in *Hdh*^*150Q/+*^ and *Hdh*^*150Q/150Q*^ mice. Subtle upregulation of p62 was observed in *Hdh*^*150Q/150Q*^. Rhes deletion has no impact on DARPP-32 or p62 levels. Rhes deletion, however, diminishes hyperactive mTORC1 signaling in *Hdh*^*150Q/150Q*^. Finally, Rhes levels are also downregulated in the striatum of HD mice in an mHTT-copy dependent manner.

## Discussion

This study reports that global Rhes deletion in a knock-in (KI) HD model provides mild benefits on selected behaviors.

The development of animal models mimicking the human HD remains a significant challenge. Transgenic (Tg) mice models that express a shorter fragment of mHTT (82 to 125 CAG repeats) show the rapid onset of motor symptoms, neuronal death, and premature demise^[63-66]^. In contrast, knock-in (KI) mice models with insertion of mHTT into exon 1 with superlong expansions (>150Q) develop slow progression, mild symptoms, or no observable HD-like symptoms^[67, 68]^. In humans, the adult-onset HD comprises >36 CAG, while the juvenile-onset >60 CAGs. The mechanisms for the paradoxical effect– while a human with >60 CAG develops severe disease and premature death while these phenotypes are absent in mouse with >150Q– remain unclear. However, the differences in the mHTT load and mHTT toxic species may determine the onset and the severity of the disease between humans and KI mice. For example, in the KI mouse models, the proteolytic mechanisms that generate toxic forms of mHTT may be suboptimal, or the KI models may have developed compensatory mechanisms that may prevent the severity of the disease.

Previous results indicate that Rhes deletion affords protections from behavioral and pathological deficits in Tg (N171-82Q and R6/2) HD animals^[45, 46]^. Here we found Rhes deletion in a milder HD model, Hdh^*150Q*^ KI mice^[47, 58]^, afforded only mild protection (Fig. 1). We found that DARPP-32, a commonly used striatal marker for striatal abnormalities, is diminished in Hdh150Q KI mice striatum (Fig. 2, 3), but these mice do not display any significant motor coordination deficits (Fig. 1). Therefore, it appears that loss of DARPP-32 is not directly linked to behavioral deficits in Hdh150Q KI mice.

Interestingly, we observed that Rhes is also diminished in the striatum of KI mice (Fig. 2, 3). The possibility that downregulation of Rhes in the KI mice may prevents the severity of HD-like symptoms cannot be ruled out. Because when Rhes is replenished in the striatum of *Rhes*^*–/–*^*;Hdh*^*150Q/150Q*^ mice, it elicited rapid motor coordination deficits^[45]^. Therefore, the downregulation of Rhes in KI striatum most likely a protective response. Nevertheless, Rhes deletion afforded protection against specific tests such as ledge, clasping, and tremor (observed at 100 weeks) (Fig. 1) as well as hyperactivation of mTORC1 signaling in *Rhes*^*–/–*^*;Hdh*^*150Q/150Q*^ (Fig. 3). Thus, residual Rhes in the striatum may still promote abnormalities in HD mice.

The mechanisms by which Rhes promotes neurodegeneration are also evolving. Rhes associates with mHTT and induces its SUMO modification, which leads to enhanced soluble forms of the mHTT^[50, 69]^. The downstream pathology may occur via more than one route. Particularly Rhes-mediated cell-to-cell transport of mHTT, which is partly SUMO-dependent, through tunneling-like nanotubes or cytoneme-like structures may elicit non-cell-autonomous toxicity^[43, 70]^.

Multiple lines of evidence link Rhes in the autophagy regulation. Rhes can activate Akt/mTORC1 signaling to inhibit autophagy^[30, 31, 71]^. Rhes binds to Beclin1 and promotes autophagy independent of mTORC1^[72]^. Rhes binds to Nix and promotes selective autophagy (mitophagy)^[44]^. Moreover, we showed that SUMO regulates mTORC1 and autophagy signaling and prevents HD-related deficits in Q175DN-KI model that show enhanced mTORC1 activity^[54, 73, 74]^. Previous pharmacological studies with mTORC1 inhibition showed beneficial effects in HD models^[54, 75]^. Thus Rhes-SUMO may orchestrate HD pathogenesis via mHTT transport as well as mTORC1/autophagy signaling.

Besides HD, Rhes is also implicated in Tau toxicity. Pathogenic tau (*MAPT)* mutations diminished Rhes expression in the hiPSC-derived neurons, and depletion of Rhes attenuated behavioral and inflammatory abnormalities in rTg4510 mice via lysosomal activation mechanism^[41]^. Histopathology evaluation of tauopathies revealed Rhes is mislocalized or diminished in the presence of abnormal tau in the entorhinal cortex, hippocampus, and superior frontal gyrus. Such specific altered neuronal distribution of Rhes is considered as a novel hallmark of all tauopathies^[42]^. Thus, alteration of Rhes localization and expression may be a common feature in neurodegenerative diseases.

In summary, our data indicate that global Rhes deletion in a less severe Hdh150Q KI model prevents only selected behavioral and biochemical deficits. Although cellular models and preclinical models could help identify new therapeutic targets, there is a growing difficulty translating these findings to humans. Recent failures of clinical trials in neurodegenerative diseases^[76]^, including HTT ASOs in HD, indicate incompatibility of murine models with clinical outcomes. Because multiple cellular and preclinical models link Rhes to both mHTT and tau toxicity, the Rhes-targeted therapy is a valuable option. Whether such treatment will diminish neuronal toxicity and increase survival can only be appreciated after the clinical trials and positive therapeutic outcomes.

## Materials and Methods

### Reagents and Antibodies

All reagents were purchased from Millipore-Sigma unless indicated otherwise. The following commercial antibodies were used: Huntingtin (WB-1:3000 clone 1HU-4C8, no. MAB2166), was obtained from Millipore-Sigma. Antibodies for DARPP-32 (WB-1:20000, no. 2306) mTOR (WB-1:3000, no. 2983), pS6K (WB-1:1000, no. 9234), S6K (WB-1;1000, no. 9202), p4EBP1 (WB-1:1000, no 9451), 4EBP1 (WB-1:15000, no. 9644), Rheb (1:2000, no. 13187), p62 (WB-1:1000, no. 39749), and LC3B (WB-1:2000, IHC-1:200, no. 3868) were from Cell Signaling Technology. Antibody against GFAP (WB-1:2000, no. 13-0300) was from ThermoFisher Scientific. Rhes antibody (WB-1:1000, RHES-101AP) was from Fabgennix. Antibodies for β-actin (WB-1:20,000; no. sc-47778), was obtained from Santa Cruz Biotechnology. HRP-conjugated secondary antibodies: goat anti-mouse (1:10,000; no.115-035-146), and goat anti-rabbit (1:10,000; no. 111-035-144) were from Jackson ImmunoResearch Inc.

### Animals

Hdh150Q knock in knock-in heterozygous (B6.129P2-*Htt*^*tm2Detl*^/150J # 004595) and C57BL/6J (JR # 000664) control mice were from The Jackson laboratory, Bar Harbor, ME. Rhes KO mice were from Alessandro Usiello were bred to produce Hdh150Q-Rhes KO and littermate groups, Hdh150Q (HD Het and HD homo), Rhes KO and WT, and the genotypes were confirmed by an established PCR protocol, as described before^[45]^. Behavior evaluation was performed as indicated before^[55]^. Number of animals used are indicated in Table 1.

### Behavioral assay

Behavioral testing was performed by the experimenter in an unbiased and blinded manner for the animal’s genotype. All behavioral testing was performed during the light phase of the light-dark cycle (8:00 a.m. and 12:00 p.m), as described before^[55]^. Rotarod output was measured on day 1, open field on day 2, and the battery of behavioral tests on day 3 of each week of behavioral testing, to avoid variability in time of testing.

### Rotarod

Every month, rotarod behavioral assay was conducted using a linear accelerating rotation paradigm (Med Associates Inc.) in three trials separated by 20 minutes. The mice were placed on the apparatus at 4 rpm and were subjected to increasing rpm, accelerating to 40 rpm over the course of a maximum of 5 min. The average of the three trials over four days was used to measure each mouse’s total latency to fall.

### Open field activity

EthoVision XT software (Noldus Information Technology) was used to measure total distance traveled and velocity of mice during a single 30-minute session in which a mouse was positioned in the center of a square enclosure and total distance traveled was measured.

### Behavioral battery

Subjective behavioral testing was adapted from a previous report^[55, 77]^. The battery of tests, each performed in triplicate, measured ledge walking, clasping, gait, kyphosis, and tremor. Individual measures were graded on a scale of 0 (no phenotype) to 3 (worst manifestation), as previously mentioned^[55, 77]^. To determine tremor, mice were placed in a clean cage and observed for 30 s. Each mouse was scored as follows: 0, no signs of tremor; 1, present but mild tremor; 2, severe intervals of tremor or constant moderate tremor; 3, outrageous chronic tremor. The mean of all five behavioral battery tests was used to create the composite ranking. Behaviors procedures were approved by the Scripps Research Institute Florida Institutional Animal Care and Use Committee.

### Western blotting

Western blotting from striatal tissue were carried as described before^[78, 79]^. Striatal tissue was rinsed briefly in PBS and directly lysed in lysis buffer [50 mM Tris-HCl (pH 7.4), 150 mM NaCl, 1% Triton X-100, 1x protease inhibitor cocktail (Roche, Sigma) and 1x phosphatase inhibitor (PhosStop, Roche, Sigma)], sonicated for 2 x 5 sec at 20% amplitude, and cleared by centrifugation for 10 min at 11,000 g at 4°C. Striatal cells or human fibroblasts were lysed in radioimmunoprecipitation assay (RIPA)buffer [50 mM Tris-HCl (pH 7.4), 150 mM NaCl, 1.0% Triton X-100, 0.5% sodium deoxycholate, 0.1% SDS] with 1x complete protease inhibitor cocktail and 1x phosphatase inhibitor, followed by a brief sonication for 2 x 5 sec at 30% amplitude and cleared by centrifugation for 10 min at 11,000g at 4°C. Protein concentration was determined with a bicinchoninic acid (BCA) protein assay reagent (Pierce). Equal amounts of protein (20-50 μg) were loaded and were separated by electrophoresis in 4 to 12% Bis-Tris Gel (ThermoFisher Scientific), transferred to polyvinylidene difluoride membranes, and probed with the indicated primary antibodies. HRP-conjugated secondary antibodies were probed to detect bound primary IgG with a chemiluminescence imager (Alpha Innotech) using enhanced chemiluminescence from WesternBright Quantum (Advansta) The band intensities were quantified with ImageJ software (NIH). Total proteins were then normalized to actin. Phosphorylated proteins to their respective total proteins.

### Statistical analysis

Data are presented as mean ± SEM as indicated. Except where stated all experiments were performed at least in biological triplicate and repeated at least twice. The mouse behavioral and the data analysis was carried out in a blinded manner. Images were quantified using ImageJ (FIJI). Behavioral data consisted of categorical and continuous outcomes. Categorical data was analyzed using Fisher’s Exact test. Statistical comparison was performed between groups using two-tailed Student’s t-test, one-way analysis of variance (ANOVA) followed by Tukey’s multiple comparison test. Significance was set at p < 0.05. All statistical tests were performed using Prism 9.0 (GraphPad software).

## Acknowledgments

We would like to thank Melissa Benilous for her administrative help and the members of the lab for their continuous support and collaborative atmosphere. We like to thank members at the Scripps proteomics and genomic core for their help and expertise. This research was supported by grant awards from NIH/NINDS R01-NS087019-01A1, NIH/NINDS R01-NS094577-01A1, a grant from Cure for Huntington Disease Research Initiative (CHDI) foundation and Scripps bridge funding.

## Author contributions

S.S conceptualized the project and co-designed experiments with J. H and N. S who performed mouse behavior. N.S performed western blotting experiments and analyzed the data. Sw. S. carried out breeding and assisted in the behavioral analysis. S.S wrote the manuscript with inputs from coauthors.

## Notes

### Competing Interest Statement

The authors have declared no competing interest.

